# Efficient meta-analysis of multivariate genome-wide association studies with Meta-MOSTest

**DOI:** 10.1101/2022.08.18.504383

**Authors:** Aihua Lin, Alexey Shadrin, Dennis van der Meer, Guy Hindley, Weiqiu Cheng, Ida Elken Sønderby, Shahram Bahrami, Kevin S O’Connell, Zillur Rahman, Nadine Parker, Olav B Smeland, Chun C. Fan, Dominic Holland, Anders M Dale, Ole A Andreassen, Oleksandr Frei

## Abstract

**Motivation:** Genome-wide association studies (GWAS) have been successful in identifying genetic variants associated with a particular phenotype. However, many complex phenotypes are influenced by multiple genetic variants with small effects. Detecting the genetic pleiotropy can provide insights into biological mechanisms influencing complex human phenotypes. The recently developed Multivariate Omnibus Statistical Test (MOSTest) has proven to be efficient and powerful, suited for complex large-scale data. The method substantially increased discovery of genetic variants associated with brain MRI phenotypes in the UK Biobank compared to conventionally use multivariate approach. Here we extend the MOSTest to meta-analysis (Meta-MOSTest), facilitating data analysis of multiple phenotypes across multiple cohorts. We evaluated our updated approach in the UK Biobank using brain MRI phenotypes, by comparing the discovery yield of the single-cohort MOSTest versus Meta-MOSTest through simulating sub-cohorts of different sample sizes from 265 to 26501 subjects.

**Results:** Our method works efficiently on large-scale cohorts with a large number of MRI phenotypes. We found that lower per-cohort sample sizes resulted in a reduced discovery yield indicating a loss of statistical power. However, with a minimum sample size of 250 subjects across cohorts, Meta-MOSTest was equivalent to MOSTest on discovery yield while maintaining a well-calibrated type I error and equivalent statistical power. We conclude that Meta-MOSTest is a useful tool for multivariate analysis across separate brain imaging genetics cohorts.

**Availability and implementation:** All codes are freely available on GitHub: <u>MOSTest</u> and <u>Meta-MOSTest</u>.

## 1 Introduction

Genome-wide association studies (GWAS) aim to identify genetic variants associated with a specific phenotype. Over the past decade, thousands of genetic variants associated with complex phenotypes have been discovered [1, 2]. These findings have provided insights into the molecular mechanisms of complex phenotypes. However, GWAS discoveries largely depend on sample size. The single nucleotide polymorphisms (SNPs) discovered by a typical GWAS only explain a small fraction of the heritability of complex phenotypes, and a large fraction of potentially relevant SNPs remain to be identified. Worldwide collaborations have increasing sample sizes. To this end, meta-analysis as a method to aggregate summary statistics across similar GWAS has been developed and applied with success [3]. The Enhancing Neuroimaging Genetics through Meta-Analysis and Psychiatric Genomics Consortium (PGC) are two examples of international collaborative consortia that combine GWAS from multiple cohorts through meta-analysis [4, 5]. They have successfully detected a series of genotype-phenotype associations that would have remained undetected by the participating cohorts alone [6, 7]. In addition, meta-analysis simplifies data sharing, keeping sensitive individual data at the cohort level and sharing only non-sensitive summary data. It also allows processing different cohorts separately with different protocols mitigating cross-cohort heterogeneity.

GWAS is a mass-univariate approach, independently testing each genetic variant for association with a given phenotype. However, there is extensive pleiotropy across complex phenotypes, defined as the same genetic variants influencing multiple different phenotypes [8]. Identifying pleiotropic variants helps to provide mechanistic insights into the underlying biology of complex human phenotypes. It can also be leveraged to boost statistical power and increase genetic discovery, compared to univariate GWAS which yield a high burden of multiple testing when applied to multiple phenotypes concurrently. For this purpose, the multivariate GWAS approaches present advantages compared to a standard univariate GWAS [9–13]. These multivariate approaches test whether the genotype vector of a SNP is associated with at least one of the multiple phenotypes. A recently developed multivariate analysis tool based on GWAS, namely Multivariate Omnibus Statistical Test (MOSTest), outperforms previously published multivariate approaches both in discovery yield and computational efficiency [14]. Leveraging the UK Biobank’s large sample of genotyped participants with high quality neuroimaging data, including phenotypes of cortical area, thickness and subcortical volumes, the method discovered 3 times more genomic loci than commonly used Min-P method [14]. The usage of genotype permutations to estimate the distribution of association test statistics under the null hypothesis enables the method to be more robust compared to analytical approximations. Further, the method was extended to analyze vertex-wise structural MRI phenotypes which yielded 8 times more loci compared to the min-P method [15], and to measures of cognition and personality with similar improvements in genetic discovery [16]. Strong replication of MOSTest-discovered loci was reported in [17] by deploying the PolyVertex Score (PVS) approach [18].

Here we present a meta-multivariate analysis tool that improves the discovery of genotype-phenotype associations by leveraging pleiotropy. It combines meta-analysis and MOSTest (Meta-MOSTest) which efficiently synthesizes data from large-scale cohorts with many phenotypes. Previous multivariate methods, like MetaCCA [19], MetaUSAT [13] and Meta-MltiSKAT[20], can only handle a small group of cohorts with up to 50 phenotypes. Our Meta-MOSTest method works efficiently on large-scale cohorts even with hundreds of phenotypes while maintaining a well-calibrated type I error. Additionally, we studied the relationship between the discovery yield and the number of cohorts included, keeping total sample size constant. We showed that for 171 regional MRI phenotypes, our updated above approximately 250 subjects. This demonstrates that the fast computation framework of Meta-MOSTest is powerful and robust enough to calculate the genetic associations for hundreds of phenotypes across multiple cohorts.

## 2 Method and Materials

Meta-MOSTest builds on the single-cohort MOSTest by extending the procedure to multiple cohorts. This involves three main steps: univariate genotype-phenotype association test within each cohort, followed by integration of univariate statistics across cohorts, and finally the MOSTest procedure. For easy illustration, we let *y_ik_* and *g_ij_* represent the *kth* phenotype vector and the *jth* genotype vector of the *h* cohort.

### Univariate genotype-phenotype association test within each cohort

After regressing out all the covariates and global features of each phenotype, we carried out the univariate association test between *y_ik_* and *g_ij_* using the standard additive linear regression model within each cohort. The statistical significance was assessed by Pearson’s correlation coefficient. The statistical significance of the univariate association between *y_ik_* and *g_ij_* was assessed by the Pearson correlation coefficient as: *r_ijk_* = *corr*(*y_ik_*, *g_ij_*). This is equivalent to testing the significance of the effect size *β_ijk_* from the linear regression, where *β_ijk_* = *r_ijk_ S_yik_*/*S_gij_*, with *S_yik_*, *S_gij_* representing the standard deviations of *y_ik_* and *g_ij_*. This is because both *β_ijk_* and *r_ijk_* are t-distributed and therefore have the same p-values.

The univariate summary statistic was calculated as 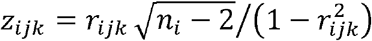, with standard error, *δ_ijk_* = *β_ijk_*/*t_ijk_*, where *n_i_* is the sample size of *h* cohort. In addition, following the same procedure, we carried out a univariate association test between each phenotype and randomly permuted genotype data producing 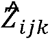. The permutation structure formed the basis of the global test statistic of the Meta-MOSTest approach.

### Integration of univariate statistics across cohorts

The global statistics of SNP *j* and phenotype *k* across cohorts *Z_jk_* were integrated by using an inverse variance based meta-analysis weighted by each cohort 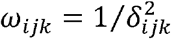, thus,

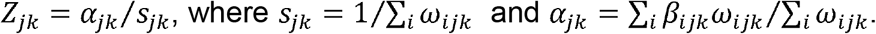

Similarly, the global statistics for the genotype permuted structure 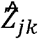 can be calculated. We prefer variance-based meta-analysis over sample-size based meta-analysis as it preserves individual properties of SNPs, such as their allele frequency – which is ignored by sample-size meta-analysis. We denote {*Z_kj_*} and 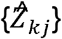 as the matrix of the global *z* scores, with rows corresponding to SNPs, and columns corresponding to phenotypes. To calculate the Meta-MOSTest test statistics, we also need the global covariance matrix *R* of *Z* scores across cohorts which is weighted by the sample size across each cohort. This is because if we let 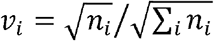 and *v_i′_* be the weights of *h* and *i*’th cohort, then

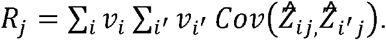

Since for independent cohorts, 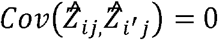, this yields 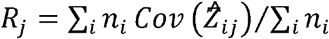. In simulations, we did not observe any inflation due to this, as any potential inflation would have been corrected by the permutation scheme.

### MOSTest procedure

The Meta-MOSTest test statistic for the *j*-th SNP is calculated by

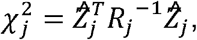

where 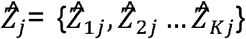 is the permuted global *Z* score vector across *K* phenotypes at the *jth* SNP. The null hypothesis of Meta-MOSTest is that the global 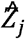 is distributed as a multi-variate normal random variable with zero mean and covariance *R_j_*. The Meta-MOSTest summary statistics are estimated by fitting the observed data under the permuted genotype scheme into a gamma distribution Γ(a, b) of shape *a* and scale *b*. The *p*-values of Meta-MOSTest are estimated from a cumulative distribution function of the gamma distribution, 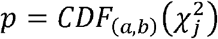 with a genome-wide significance threshold of *α* = 5 × 10^−8^.

### Locus definition

The definition of genetic loci from Meta-MOSTest is in line with the protocol implemented in FUMA [21], with the following steps:

1. A list of independent significant genetic variants is selected with genome-wide significance threshold lower than 5 × 10^−8^ and linkage disequilibrium (LD) [22] r2< 0.6 with each other.
2. Lead genetic variants are defined as those independent significant variants with LD r2< 0.1 with each other.
3. Candidate variants are those which have LD r2≥ 0.6 between them and an independent significant variant
4. The genomic locus of a lead variant are defined as min/max positional coordinates over all corresponding candidate variants.
5. Loci are merged if they are separated by less than 250kb.

LD correlations are computed from EUR population of the 1000 genomes Phase 3 data.

### Sample

To compare the discovery yield between Meta-MOSTest of multiple cohorts and MOSTest applied to data from a single cohort, we used the same sample as in the previous study [14]. T1 MRI scan data were obtained from the UK Biobank released up to April 2019 under accession number 27412. Details about the data composition and gathering protocols were described in [23]. Participants were selected from White European ancestry that had undergone the neuroimaging protocol and passed genetic quality control procedures described below. This provided the sample size of our analysis *n*◻=◻26,501, with a mean age of 64.4 years (SD◻=◻7.5) and 51.7% were female.

### Data Preprocessing

T1-weighted scans were collected from three identically configured Siemens Skyra 3T scanners from the United Kingdom, more details can be found in [23]. We process the data with the standard “recon-all-all” processing pipeline of Freesurfer v5.3 and extracted the sets of regional subcortical and cortical morphology phenotypes, as well as estimated intracranial volume (eICV). Both left and right hemisphere phenotypes were included in this study.

To set up a multi-cohort scenario, we distributed the participants from three scanners into cohorts of similar sample sizes to get an overview of the relationship between sample size and discovery yield. To make sure we are studying regional rather than global effects, we regressed out age, sex, scanner site, Euler number, and the first 20 genetic principal components within each cohort. We further regressed out a global phenotype specific to each of the feature subsets: eICV for the subcortical volumes, mean thickness for the regional thickness phenotypes, and total surface area for the regional surface area phenotypes. This resulted in 68 cortical surface area and thickness phenotypes and 35 subcortical volume phenotypes (*Supplementray Table 1*). To achieve a normally distributed input for the association test between genotype and phenotype within each cohort, a rank-based inverse-normal transformation to the residuals of each phenotype was performed. We performed the same preprocessing for the simulation analysis as in the primary analysis.

We used the UK Biobank v3 imputed data as our genotype data [24]. In addition, we filtered out individuals with more than 10% missing rate, genetic variants with more than 5% missing rate, variants failing the Hardy–Weinberg equilibrium test at p◻=◻1◻×◻10−9 and variants with a minor allele frequency below 0.005, leaving 7,428,630 variants.

### Analytical Design

We applied our Meta-MOSTest approach to MRI regional morphology phenotypes from the UK Biobank with a total sample size of 26,501, including 68 regional cortical area phenotypes, 68 regional cortical thickness phenotypes, 35 subcortical volumes and a combination of all 171 features (‘all phenotypes’). To compare the discovery yield between MOSTest and Meta-MOSTest and to investigate the relationship between sample size and discovery yield, we randomly distributed a total of 26,501 subjects into a different number of cohorts with similar sample sizes. The sample sizes were defined by the usual used range of sample sizes included in PGC and ENIGMA studies [10], with a per-cohort sample size of 26501, 5300, 1050, 530, 265, 133 and 66.

## 3 Results

### Genetic variant discoveries

*Fig. 1* shows quantile-quantile (Q-Q) plots of analysis on ‘all phenotypes’ at a significance threshold of *α* = 5 × 10^−8^ where subjects were respectively distributed into 1, 5, 25, 50,100, 200 and 400 cohorts for each meta-analysis scenario, corresponding to per-cohort sample sizes of 26501, 5300, 1050, 530, 265, 133 and 66. Further decreases in the number of power. However, there was no significant reduction in the enrichment of statistical association when the per-cohort sample size was decreased from 26501 to 265. This feature was also present in Q-Q plots of the other phenotypes (S*upplementary fig. 1-3*). *Table 1* and Manhattan plots (S*upplementary fig. 4-7*) showed that further decreased in the number of per-cohort sample size from 265 to 66, resulted in a reduced discovery yield, from 281 to 81 for the ‘all phenotypes’, indicating a loss of statistical power, but the identity of the most significant lead SNPs was highly consistent. For the analysis of ‘all phenotypes’ with a per-cohort sample size 66 (400 cohorts), 81 significant genetic variants were identified, more than the commonly used Min-P method, where 53 loci were reported [14].

**Figure 1.**
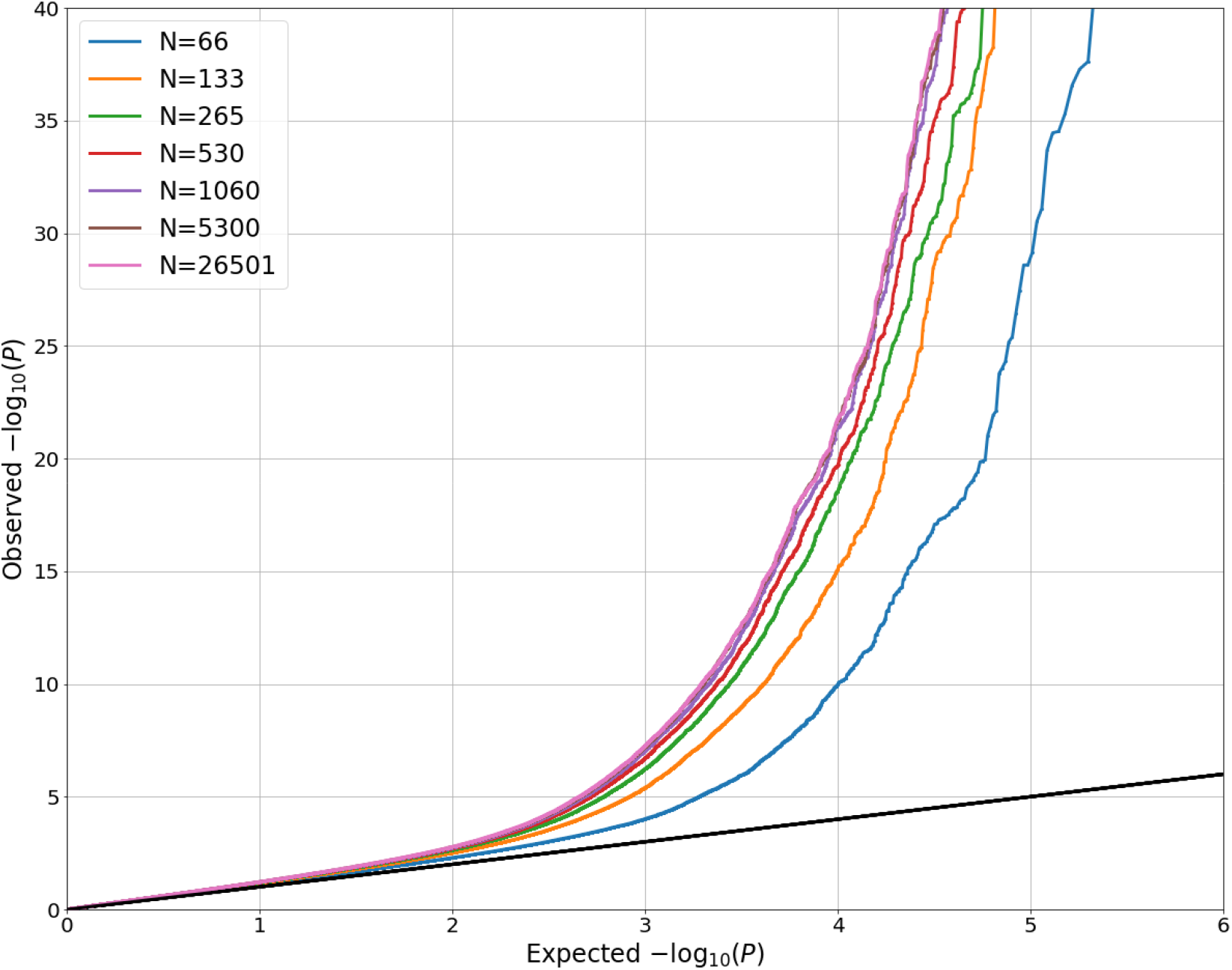
QQ plots of “all phenotypes” with different per-cohort sample sizes (N=26501, 5300, 1060, 530, 265, 133 and 66 corresponding to the number of cohorts of 1, 5, 25, 50, 100, 200 and 400.

**Table 1.** Number of significant genetic variants on analysis of cortical area, cortical thickness, subcortical volume and ‘all phenotypes’ at significance threshold of with different per-cohort sample sizes.

| Number of cohorts | Per-cohort sample size | Number of significant variants |  |  |  |
| --- | --- | --- | --- | --- | --- |
|  |  | cortical area | cortical thickness | subcortical volume | ‘all phenotypes’ |
| 1 | 26501 | 136 | 71 | 175 | 342 |
| 5 | 5300 | 136 | 72 | 176 | 332 |
| 25 | 1060 | 135 | 63 | 170 | 336 |
| 50 | 530 | 128 | 63 | 161 | 312 |
| 100 | 265 | 113 | 45 | 147 | 281 |
| 200 | 133 | 94 | 42 | 101 | 214 |
| 400 | 66 | 1 | 15 | 49 | 81 |

### Validation

To test our Meta-MOSTest approach, we simulated a scenario with pre-defined causal genetic variants. We used the same genotype data as in the main analysis and randomly selected 1000 variants on chromosome 21 as the causal variants (β≠0). Then, we generated 10 quantitative phenotypes with heritability 0.04 using the additive genetic model y = *Gβ* + *ε*, where G is the genotype matrix and *ε* is a normally distributed residual. For simplicity, all phenotypes shared the same group of causal variants but with different effect *β*. Thus, the type I error, α was calculated as the proportion of the significant variants among all variants with null genetic effects. We observed that type I error matched the pre-selected threshold(*α* = 0.0001, 0.01 and 0.05) for all scenarios with different per-cohort sample sizes (S*upplementary fig. 8*).

The statistical power was calculated from the type II error, which was the proportion of non-significant variants among causal variants. It showed that the statistical power increased with higher significance threshold. At a fixed significance threshold, the statistical power decreased slightly with smaller per-cohort sample sizes (S*upplementary fig. 9*).

## 4 Discussion

The main finding of the present study was that the multivariate GWAS analytical tool Meta-MOSTest improves discovery of SNPs with distributed effects across brain MRI phenotypes in a meta-analysis of multiple cohorts, with a similar performance as MOSTest in single cohort analysis when the per-cohort sample size was above 250 subjects [14]. This demonstrates that the fast computation framework of Meta-MOSTest is powerful and robustly identifies the genotype-phenotype associations for hundreds of phenotypes across multiple cohorts. In addition, meta-analysis provides a more reliable setup when there is strong heterogeneity across cohorts. This opens the possibility to apply a multivariate GWAS approach through meta-analysis (Meta-MOSTest) of small studies with sensitive data to boost genetic discoveries in collaborations comprising many independent samples.

We systematically investigated the relationship between discovery yield and per-cohort sample sizes for a joint meta-multivariate analysis. Meta-MOSTest includes a univariate association test between the phenotype and randomly permuted genotype data within each cohort. This is different from other existing meta-multivariate methods (MetaCCA [19], frameworks [13, 19, 20]. Thus, for the global statistic summary, we employed variance-based meta-analysis which preserved individual properties of SNPs, such as their allele frequency, which is ignored by sample-size meta-analysis. Our variance-based meta-analysis strategy also greatly reduces the computing time compared to the sample-size meta-analysis procedure. The current results provided some indications of the minimal reasonable sample size for a Meta-MOSTest analysis in imaging genetic studies. For the analysis of 171 regional MRI phenotypes with a pooled sample size of 26,511, the discovery yield decreased significantly when the per-cohort sample size was smaller than 265. Thus, it is important to have a minimum sample size for the cohorts included in imaging genetics multisite studies. This is in line with the usual used minimal sample sizes in GWAS from other meta-analysis. For example, in study of identifying the genetic variants related with cortical structure, 39 of 60 cohorts from ENIGMA have more than 265 subjects [10]. In a posttraumatic stress disorder (PTSD) genome-wide association study, 37 of 47 cohorts from PGC have sample sizes bigger than 265 [7].

There is a need for fast computation framework to handle the highly dimensional genotype-phenotype data in large, deeply phenotyped cohorts which are becoming increasingly prevalent. In contrast, other multivariate methods, including those listed above, can only handle a small group of cohorts with up to 50 phenotypes. To the best of our knowledge, Meta-MOSTest is therefore the only meta-multivariate tool that can efficiently work on large-scale genotyped cohorts with many phenotypes.

The first step of the Meta-MOSTest procedure, namely univariate association tests between the phenotypes both with the genotype data and with the permuted genotype data, can be done on each site due to data protection. Thus, distribution of Meta-MOSTest software package is necessary for performing local analysis. The results of this step do not contain individual level data and can be aggregated in a central location for the last two steps of the Meta-MOSTest procedure, i.e., integration of univariate statistics across cohorts and the Meta-MOSTest test statistics.

The current work was illustrated in a framework using neuroimaging data where all cohorts have the same sample sizes. This approach was taken because different distribution of subjects across cohorts may have different effects on the discovery yield. One of our goals is to investigate the difference of discovery yield affected by the smallest sample size, thus, the same sample size across cohort was chosen. For practical use, we will update the approach for applications with varying sample sizes across cohorts. This will be very useful for neuroimaging studies in the large ENIGMA network which have successfully applied distributed analysis to gain insight into the genetic architecture of brain structure phenotypes [4]. However, the number of brain MRI phenotypes has so far been limited. Importantly, the Meta-MOSTest approach can be applied to large-scale genotyped cohorts with a large number of phenotypes. Thus, it can also uncover more of the genetic architecture of a range of other complex human phenotypes, from mental to cardiometabolic traits. Another limitation of the current work is that the simulations were on UK Biobank data which has high homogeneity. One potential direction of our future work is to investigate the discovery yield difference when different ancestries are included.

To conclude, the Meta-MOSTest method works efficiently on large cohorts of complex phenotypes, while maintaining a well-calibrated type I error. The Meta-MOSTest approach achieved similar discovery yield as MOSTest when the per-cohort sample size was above 250 subjects. The fast computation framework of Meta-MOSTest can be used to improve genetic discovery for hundreds of phenotypes across multiple cohorts in compliance with data protection regulations.

## Data availability

The data used in the primary analysis were gathered from the public UK Biobank resource and will be made publicly available together with the code used to generate the data through the UK Biobank Returns Catalogue (https://biobank.ndph.ox.ac.uk/showcase/docs.cgi?id=1).

## Funding

This work was supported by the Research Council of Norway [#223273, #273291, #276082], KG Jebsen Stiftelsen (SKGJ-MED-021) and EU’s H2020 RIA grant #847776 CoMorMent, and SouthEast Norway Regional Health Authority (#2020060).

This work was performed on Services for sensitive data (TSD), University of Oslo, Norway, with resources provided by UNINETT Sigma2 - the National Infrastructure for High Performance Computing and Data Storage in Norway.

This research has been conducted using data from UK Biobank, a major biomedical database (www.ukbiobank.ac.uk).

