## Supplementary figures for "Efficient meta-analysis of multivariate genome-wide association studies with Meta-MOSTest"

**Supplementary Table 1.**
Regional brain morphology outcome measures, per subset, included in the study.

*Subcortical Volumes	Surface Area & Thickness*
1. Lateral Ventricle	1. Bankssts
2. Inferior Lateral Ventricle	2. Caudalanteriorcingulate
3. Cerebellum White Matter	3. Caudalmiddlefrontal
4. Cerebellum Cortex	4. Cuneus
5. Thalamus Proper	5. Entorhinal
6. Caudate	6. Fusiform
7. Putamen	7. Inferiorparietal
8. Pallidum	8. Inferiortemporal
9. 3rd Ventricle*	9. Isthmuscingulate
10. 4th Ventricle*	10. Lateraloccipital
11. Brain Stem*	11. Lateralorbitofrontal
12. Hippocampus	12. Lingual
13. Amygdala	13. Medialorbitofrontal
14. Accumbens Area	14. Middletemporal
15. Ventral Diencephalon	15. Parahippocampal
16. Choroid Plexus	16. Paracentral
17. 5th Ventricle*	17. Parsopercularis
18. CC Posterior*	18. Parsorbitalis
19. CC Mid Posterior*	19. Parstriangularis
20. CC Central*	20. Pericalcarine
21. CC Mid Anterior*	21. Postcentral
22. CC Anterior*	22. Posteriorcingulate

23. Precentral
24. Precuneus
25. Rostralanteriorcingulate
26. Rostralmiddlefrontal
27. Superiorfrontal
28. Superiorparietal
29. Superiortemporal
30. Supramarginal
31. Frontalpole
32. Temporalpole
33. Transversetemporal
34. Insula

Note: * not bilateral

**Supplementary figures**

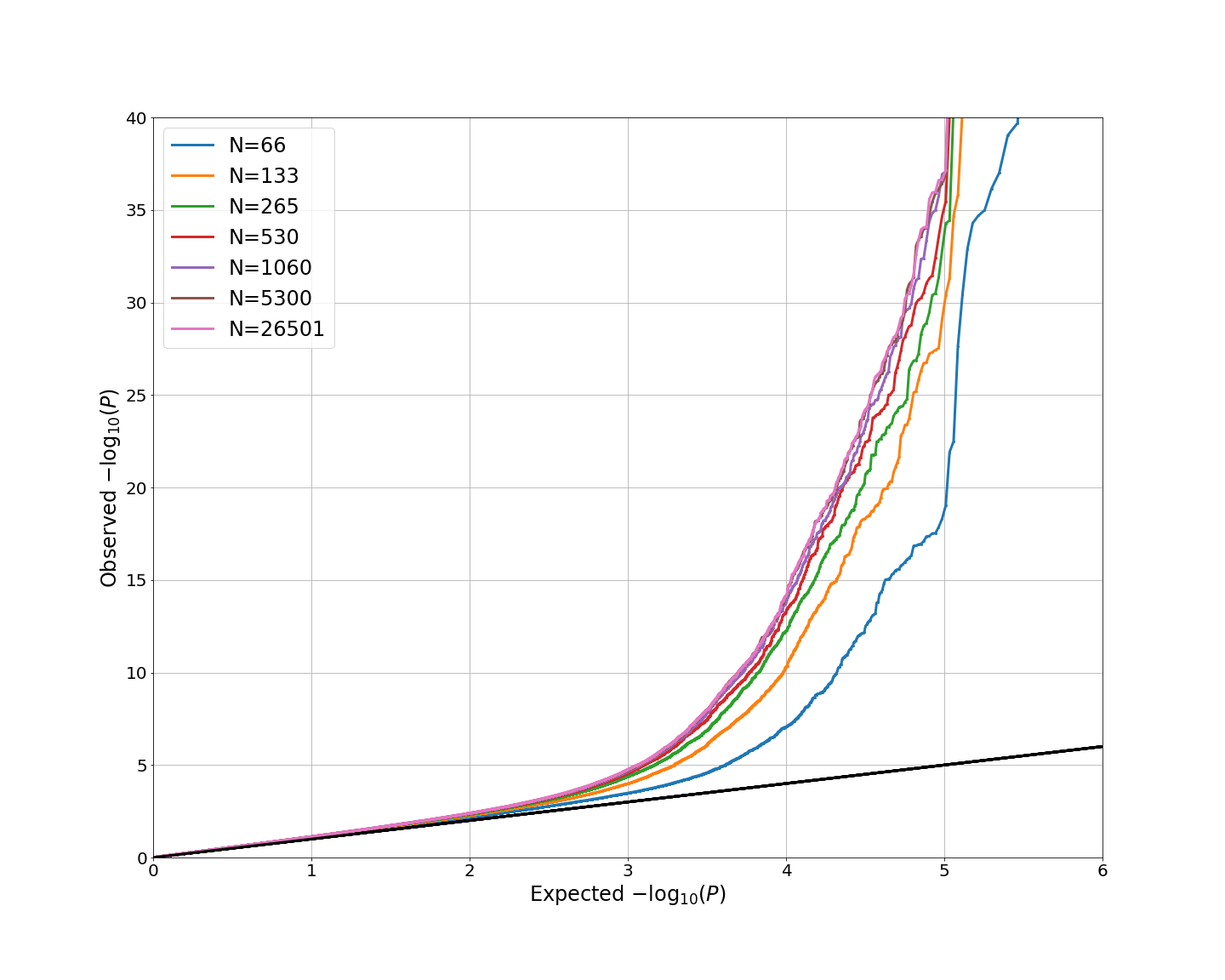

Figure 1. QQ plots of cortical area with different per-cohort sample size of each cohort (N=26501, 5300, 1060, 530, 265, 133 and 66 corresponding to number of cohorts of 1, 5, 25, 50, 100, 200 and 400) at significance threshold of $5\times{10}^{-8}$.

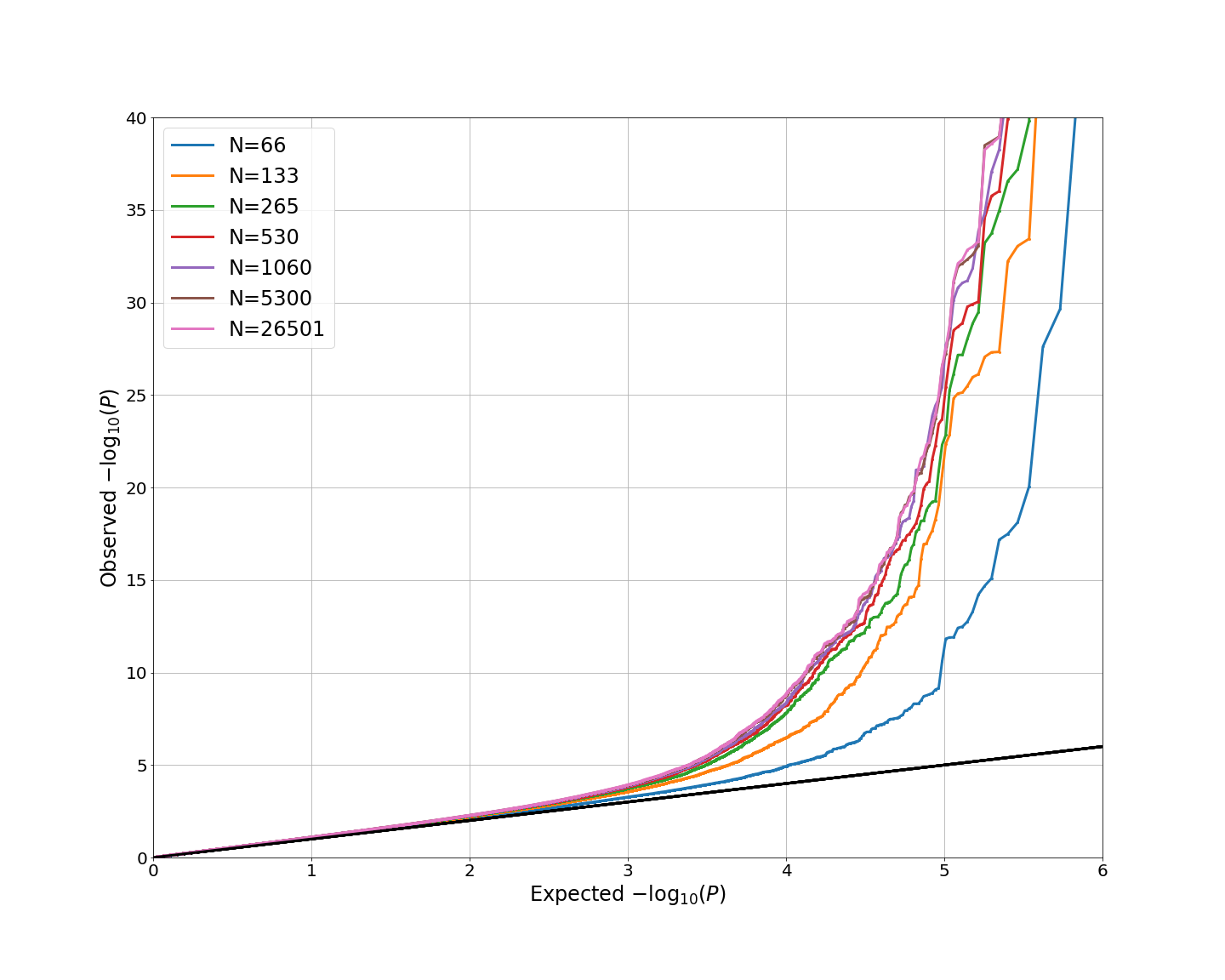

Figure 2. QQ plots of cortical thickness with different per-cohort sample size of each cohort (N=26501, 5300, 1060, 530, 265, 133 and 66 corresponding to number of cohorts of 1, 5, 25, 50, 100, 200 and 400) at significance threshold of $5\times{10}^{-8}$.

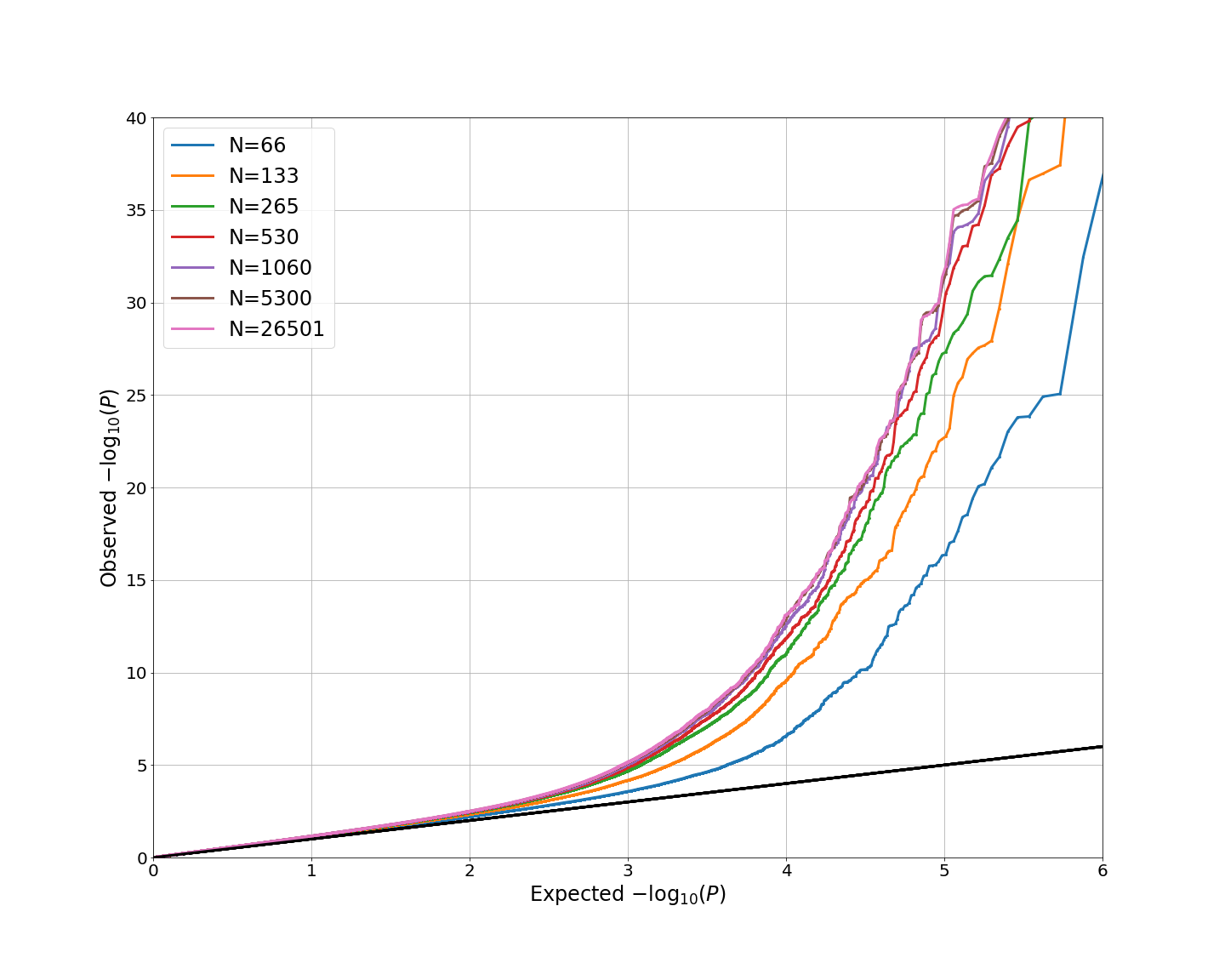

Figure 3. QQ plots of subcortical volume with different per-cohort sample size of each cohort (N=26501, 5300, 1060, 530, 265, 133 and 66 corresponding to number of cohorts of 1, 5, 25, 50, 100, 200 and 400) at significance threshold of $5\times{10}^{-8}$.

(a)

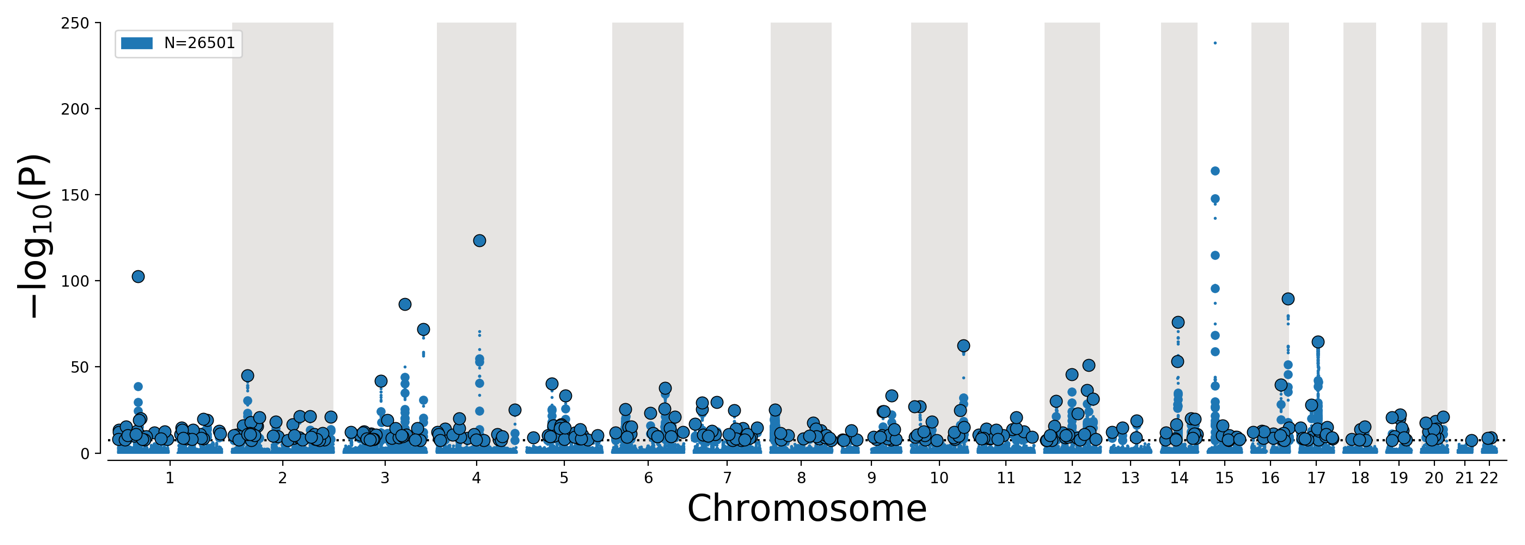

(b)

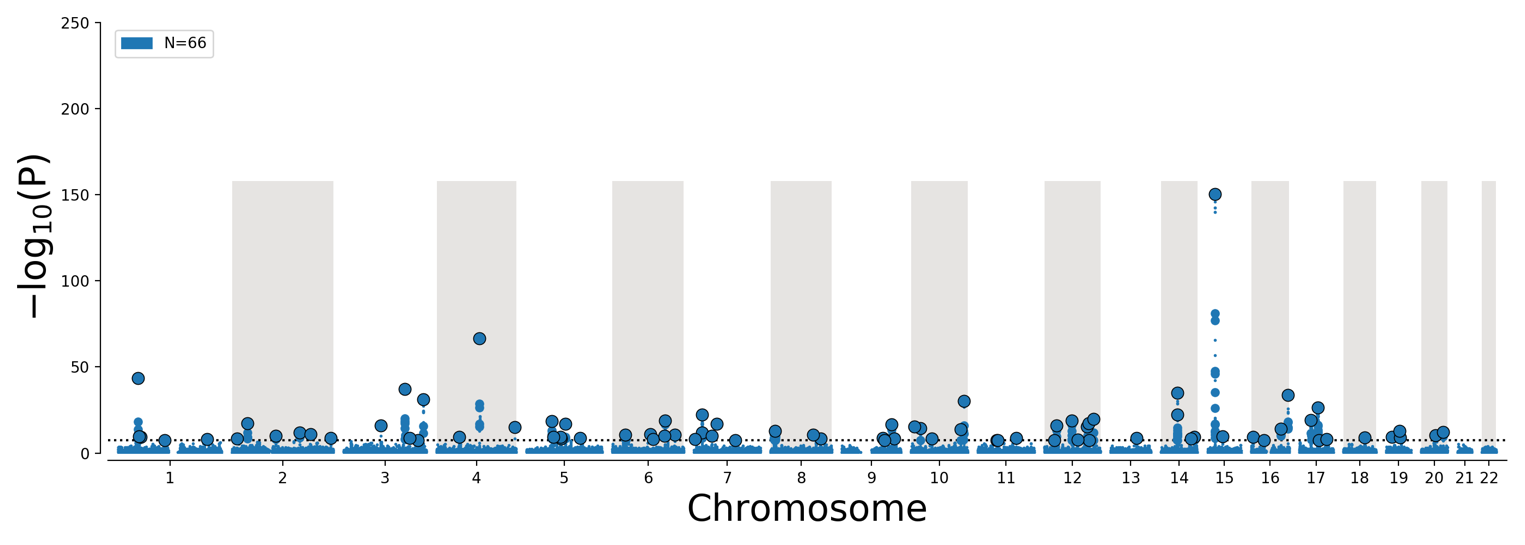

Figure 4. Manhattan plots of “all phenotypes” with (a) single cohort MOSTest and (b) Meta-MOSTest (400 cohorts of per-cohort sample size of 66).

(a)

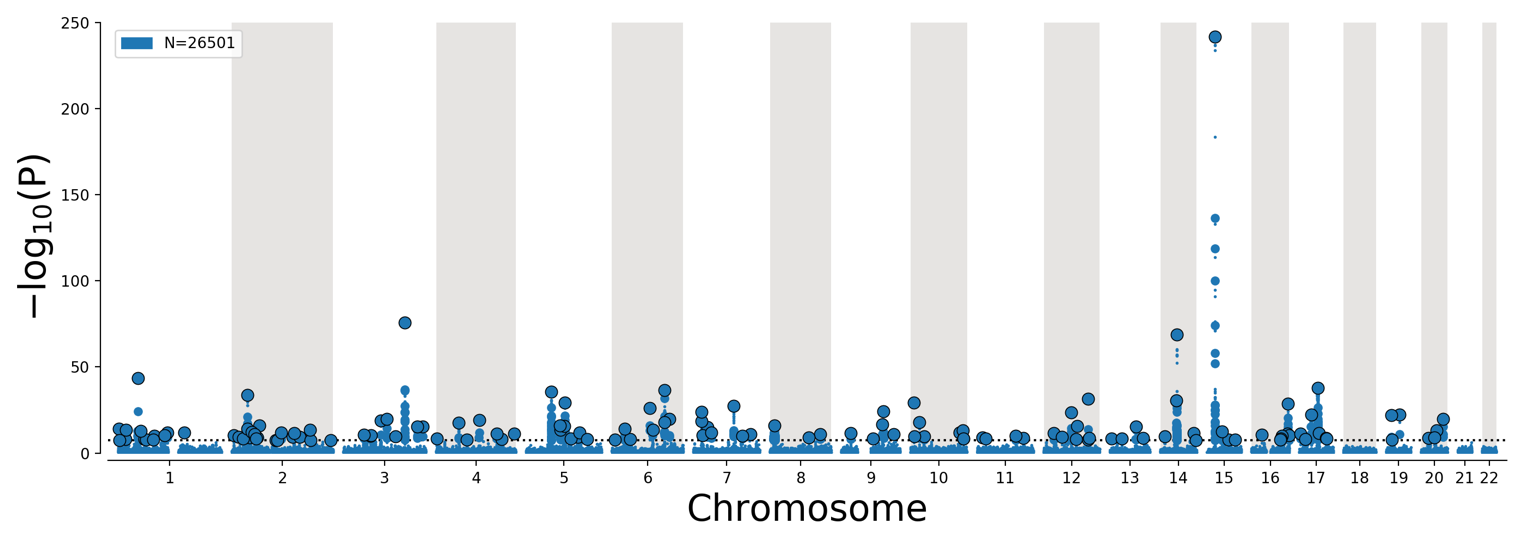

(b)

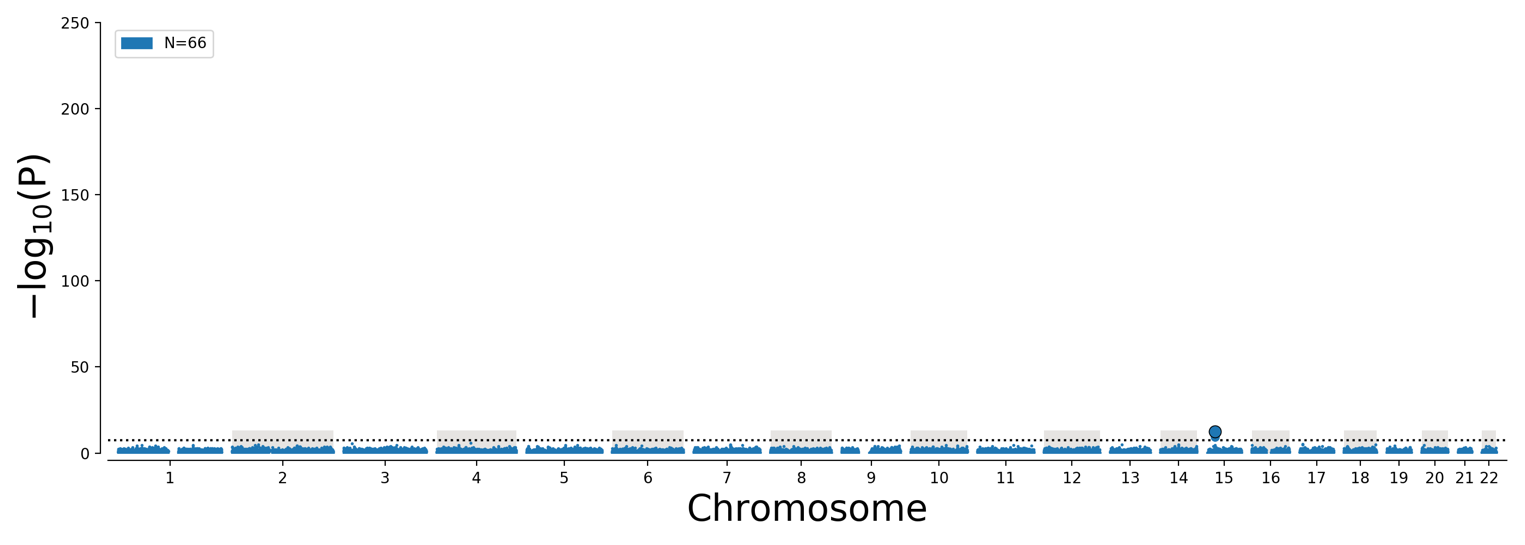

Figure 5. Manhattan plots of cortical area with (a) single cohort MOSTest and (b) Meta-MOSTest (400 cohorts of per-cohort sample size of 66).

(a)

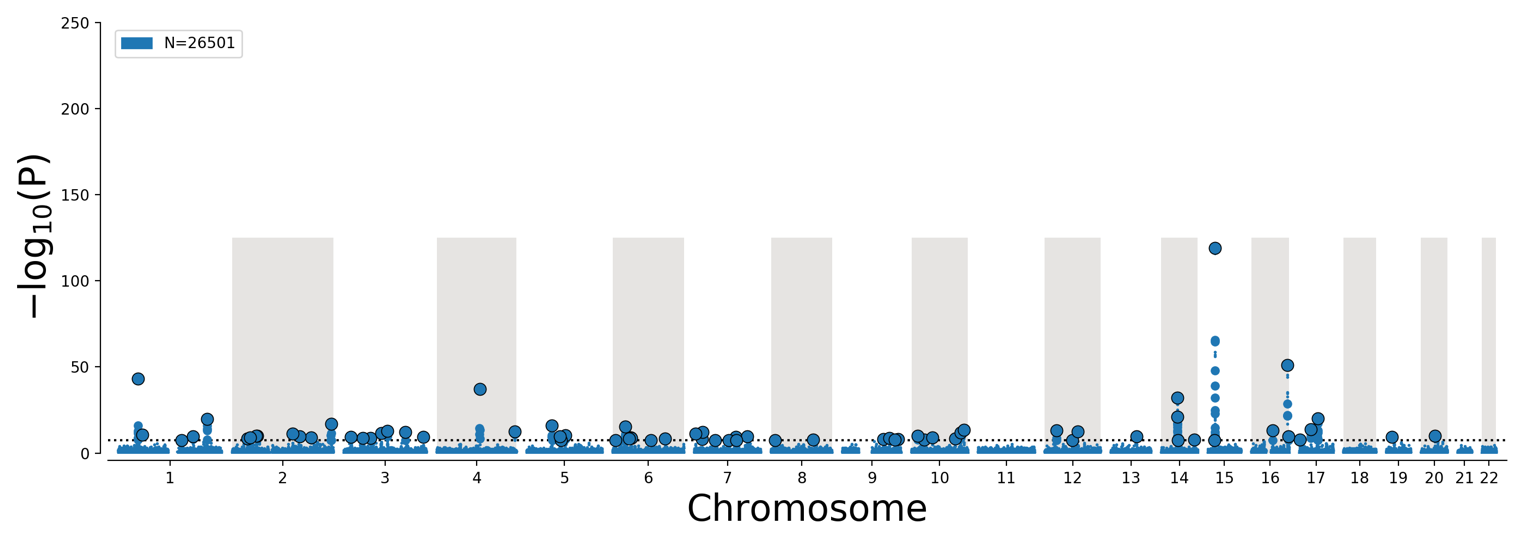

(b)

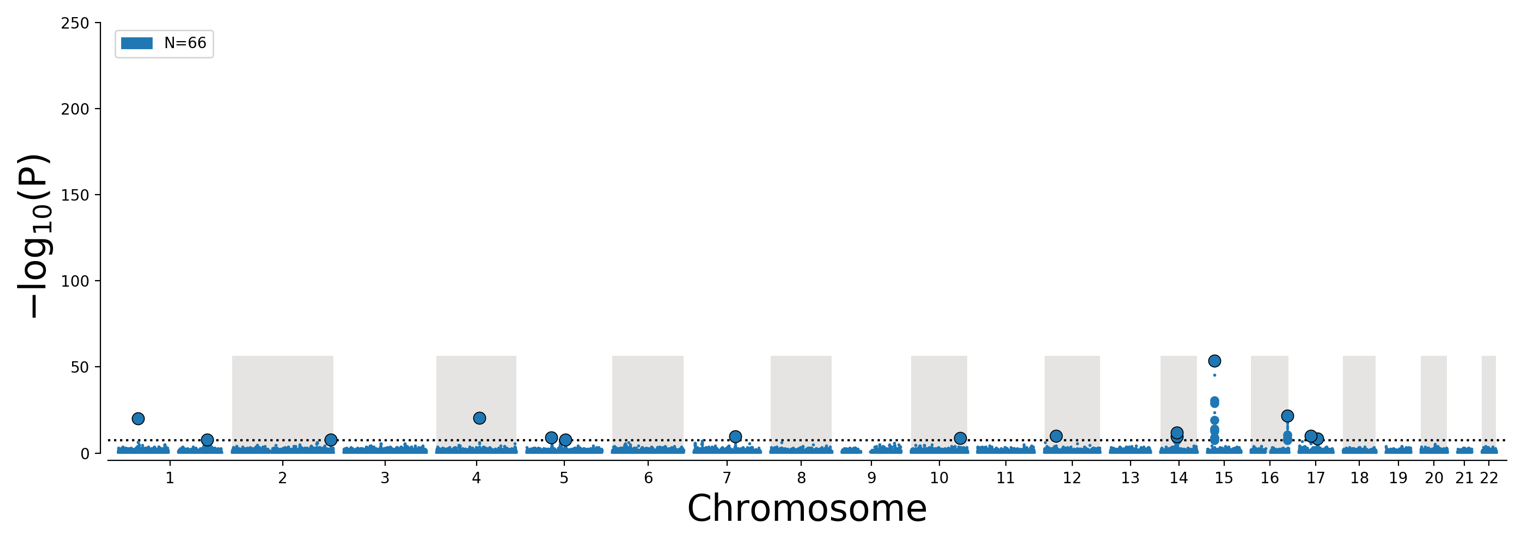

Figure 6. Manhattan plots of cortical thickness with (a) single cohort MOSTest and (b) Meta-MOSTest (400 cohorts of per-cohort sample size of 66).

(a)

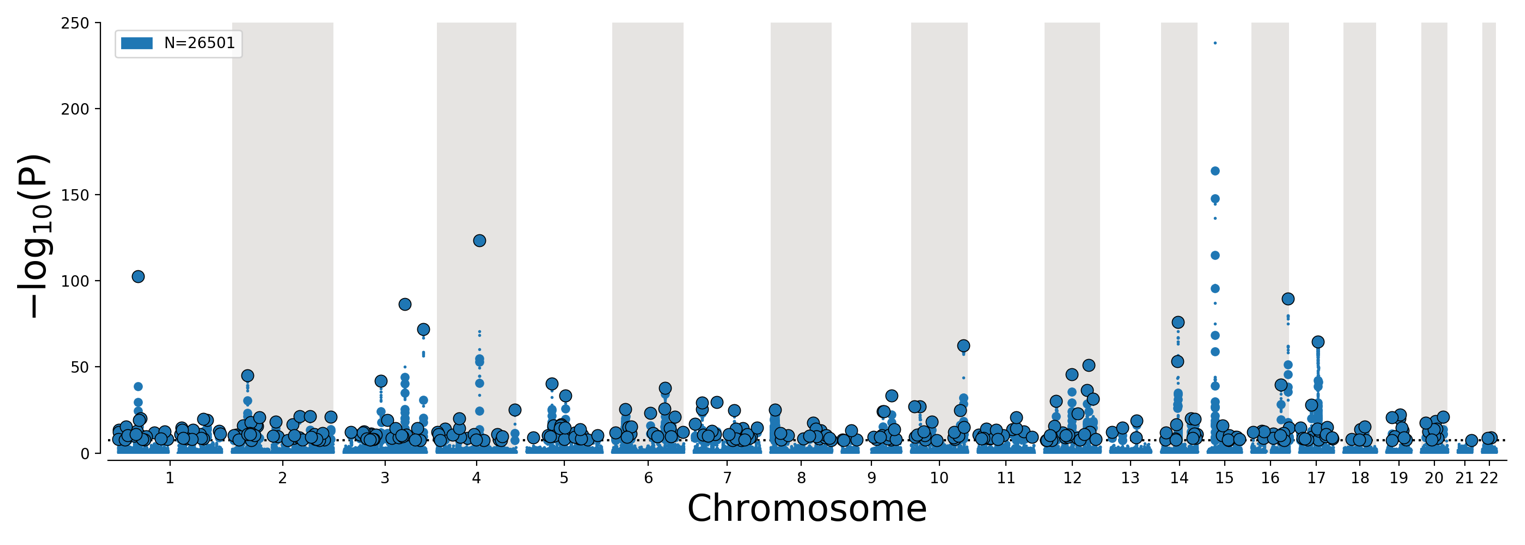

(b)

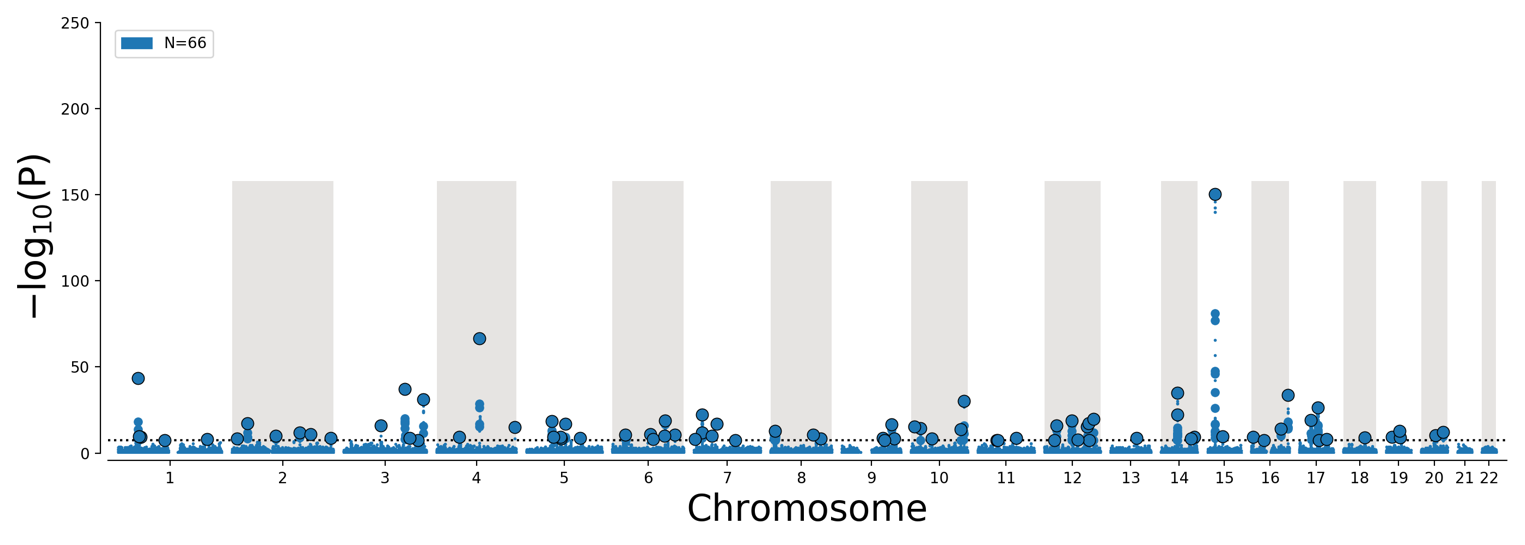

Figure 7. Manhattan plots of subcortical volume with (a) single cohort MOSTest and (b) Meta-MOSTest (400 cohorts of per-cohort sample size of 66).
